# Instantaneous polarized light imaging reveals activity dependent structural changes of dendrites in mouse hippocampal slices

**DOI:** 10.1101/523571

**Authors:** Maki Koike-Tani, Takashi Tominaga, Rudolf Oldenbourg, Tomomi Tani

**Affiliations:** Eugene Bell Center for Regenerative Biology and Tissue Engineering, Marine Biological Laboratory, Woods Hole, Massachusetts, U.S.A.; Department of Pharmacology, Tokushima-Bunri University, Kagawa, Japan.

**Keywords:** polarized light imaging, synapses, hippocampus neurons

## Abstract

Intrinsic Optical Signal (IOS) imaging has been widely used to map patterns of brain activity *in vivo* in a label-free manner. Traditional IOS refers to changes in light transmission, absorption, and scattering, which have been correlated with neuronal swelling and volume changes in the observed tissue. Here we use polarized light for IOS imaging to monitor structural changes of cellular and sub-cellular architectures of neurons due to their synaptic activity in isolated brain slices. In order to reveal fast spatio-temporal changes of birefringence associated with neuronal activity, we developed the instantaneous PolScope. The instantaneous PolScope records changes in transmission, birefringence, and slow axis orientation in tissue at high spatial and temporal resolution using a single camera exposure. These capabilities enabled us to correlate polarization-sensitive IOS with traditional IOS on the same preparations. We detected reproducible spatio-temporal changes in both IOSs at the stratum radiatum in mouse hippocampal slices evoked by Schaffer collateral stimulation in the CA1 area. Upon stimulation, changes in traditional IOS signals were broadly similar across the area, while birefringence imaging revealed local variations not seen in traditional IOS. Locations with high resting birefringence produced larger stimulation-evoked birefringence changes than those with low resting birefringence. Local application of glutamate to the synaptic region in CA1 induced increase in both transmittance and birefringence signals. Blocking synaptic transmission with CNQX and D-APV (inhibitors of AMPA-and NMDA-type ionotropic glutamate receptors, respectively) reduced the peak amplitude of the optical signals. Changes in both IOSs were enhanced by an inhibitor of the membranous glutamate transporter, DL-TBOA. Our results indicate that birefringence imaging can monitor structural alterations of dendrites subjected to excitatory synaptic transmission also associated with neuronal activity in the brain.

## Introduction

Intrinsic Optical Signal (IOS) imaging has been used as one of the most promising label-free approaches that map the patterns of brain activity *in vivo* (1–4). Traditional IOS refers to changes in optical transmission, scattering and reflectance of the tissue. These optical changes have been associated with alterations in the balance of oxy- and deoxy-hemoglobins (1, 2), swelling of neurons (5–7), changes in ionic metabolism in astrocytes (8, 9), and flavin metabolisms (10–12) that are correlated with neuronal activity *in vivo*. In isolated brain slice preparations, neuronal swelling is thought to be the major cause of IOS in response to electrical stimulations of neurons (5–7, 9). Past studies indicated that morphological changes of organelles, such as mitochondria (8, 13, 14) and intracellular vesicles (15) contribute to traditional IOS changes in primary cultured neurons and neuronal tissues.

Polarized light microscopy has been a powerful tool to investigate the dynamics of anisotropic structural signatures of molecular assemblies in living cells in a label-free and non-invasive manner. The biological polarized light microscope was developed more than a half century ago to observe the assembly/disassembly of protein polymers that produce mechanical forces for segregating chromosomes (16, 17). Later, an advanced polarizing microscope, the LC-PolScope, was built based on the traditional polarizing microscope, introducing two essential modifications: the specimen is illuminated with nearly circularly polarized light; and the traditional compensator is replaced by a liquid crystal based universal compensator (18). The LC-PolScope captures images of a specimen that is sequentially illuminated with light of five different polarization states. The raw images are then used to compute the absolute orientation of the slow axis of birefringence and the amount of differential retardation, also called retardance, at each resolved specimen point, regardless of the orientation of the object itself. Like with the traditional polarizing microscope, the LC-PolScope enables the observation of ordered structures such as the cytoskeleton and lipid membranes, without treating cells with exogenous dyes or fluorescent labels (19–21), albeit at a much higher sensitivity. By taking advantage of the high sensitivity and quantitative nature of LC-PolScope images, we have shown the anatomical structures composed of both myelinated and non-myelinated axons and bundle of dendrites, and their developmental changes in acute slice preparations of chick cerebellum (21).

In spite of its usefulness and reliability in quantitatively mapping quasi-static birefringent structures in living cells, the image acquisition time of the LC-PolScope limits the temporal resolution to about 1 s or longer. This time resolution is not sufficient to observe temporal birefringence changes in neuronal tissues subject to neuronal activity. Therefore, we developed the instantaneous PolScope, that records the optical anisotropy of observed objects using a 4-way polarization beam splitter in the imaging optics. The setup was first implemented for the instantaneous FluoPolScope developed in our laboratory (22). The time resolution of the instantaneous PolScope is as fast as 100 Hz, enabling us to monitor synaptically evoked changes in birefringence and transmittance of the stratum radiatum of area CA1 in mouse hippocampal slices. By monitoring both, changes in transmittance and birefringence, we were able to reveal cellular-or subcellular level synaptically evoked changes in neuronal structures which cannot be observed with traditional IOS alone.

## Materials and Methods

### Ethical approval

All experiments were performed according to the guidelines of the Institutional Animal Care and Use Committee (IACUC) approved by the Marine Biological Laboratory. Animal experiments were performed on brain tissue from male and female CD-1 mice (Charles River, Boston, MA) of postnatal day 4 to 20 weeks of age, housed in the animal facility at the Marine Biological Laboratory. Mice were anaesthetized using isoflurane, and then quickly decapitated.

### Preparation of brain slices and solutions

After the decapitation, the isolated brain was quickly transferred on ice-cold cutting solution containing (in mM): 205.4 sucrose, 2.5 KCl, 26 NaHCO_3_, 1.25 NaH_2_PO_4_, 0.4 CaCl_2_, 4 MgCl_2_, 10 glucose, (pH 7.4 when bubbled with a moistened mixture of 5% CO_2_ and 95% O_2_). After cooling the brain on ice for 5 min, the hippocampus together with surrounding cortex was sliced in 250 µm thick slices using a linear slicer (Dosaka Pro-7, Kyoto, Japan). Each slice was transferred onto a membrane filter (Millipore, 0.45 µm) after a short incubation in artificial cerebrospinal fluid (ACSF) containing (in mM): 124 NaCl, 2.5 KCl, 26 NaHCO_3_, 1.25 NaH_2_PO_4_, 2 CaCl_2_, 1 MgCl_2_, 10 glucose, (pH 7.4 when bubbled with a moistened mixture of 5% CO_2_ and 95% O_2_). Brain slices placed on membrane filters were incubated in a moist chamber at room temperature (22-26°C) until imaging. CNQX, D-APV, bicuculline, DL-TBOA, Dihydrokainic acid and TTX were purchased from Tocris (Bio-Techne Corporation, Minneapolis, MN).

### Primary culture

Cultured hippocampal neurons were prepared from postnatal 1-3 days of C57BL/6 mice. After anesthetizing by inhalation of isoflurane, the pups were quickly decapitated. Hippocampi were dissected out and maintained in Earle’s Balanced Salt Solution (EBSS) containing papain for 30 min at 37°C. The hippocampi were transferred to the culture media contained Neurobasal A medium, B27 supplementand 5 % fetal bovine serum (ThermoFisher Scientific, Waltham, MA), and then gently dissociated using a fire polished glass pipette. The dissociated cells were plated on poly-D-lysine coated coverslips and maintained in culture media at 37 °C in a humidified 5 % CO_2_ incubator. Twice a week, half the culture medium was replaced with new medium. The cultured neurons were used for imaging between 7 to 14 days after plating.

### Preparation setting and nerve stimulation

The brain slices were gently transferred into the imaging chamber under the dissecting microscope. During imaging the slices were continuously perfused with ACSF at a rate of 1 ml per min. ACSF was continuously bubbled with mixture of 5% CO_2_/ 95% O_2_ gas and warmed to 35°C with an in-line temperature controlling devise (Warner Instruments, Hamden, CT). An ACSF-filled glass pipette (opening diameter 10-30 µm, 1B120F-4, World Precision Instruments, Sarasota, FL) was used for Schaffer collateral nerve stimulations. The stimulus train (250 µA, 1 ms pulse duration,, 50 pulses, 40 Hz) was generated by a pulse generator (B&K Precision Corporation, Yorba Linda, CA) through an isolator (World Precision Instruments, Sarasota, FL) to induce intrinsic optical signal changes.

### LC-PolScope imaging

The principles of LC-PolScope imaging were described elsewhere (23). The brain slice was illuminated with a direct current-stabilized 100W tungsten halogen lamp through a band-pass filter (529/24 nm, Semrock Inc, Rochester, NY). The acute hippocampal slices were imaged with 20× Olympus water-dipping lens (Olympus XLUMPLANFL 20×, NA0.95). The universal compensator in the optical path of the LC-PolScope includes two liquid crystal variable retarders and a polarizer. Several predetermined polarization settings can be registered in the controller of the universal compensator and are used to sequentially acquire five raw images on a CCD camera (QImaging, Retiga 2000RV, Burnaby, Canada). The captured images were used for computing the amount of retardance, and the absolute slow axis orientation of the birefringence at each resolved specimen point. For image computation and analysis we used the OpenPolScope plug-ins for the open source image analysis platforms ImageJ and Micro-Manager (23).

### Instantaneous PolScope imaging

The instantaneous PolScope is a modification of the instantaneous FluoPolscope that we developed for monitoring the position and orientation of single emission dipoles of fluorophores in living cells (22). For the instantaneous PolScope, we use the same polarizing beam splitters in the imaging path as described in (22). For illumination, however, we use the transmitted light path, providing incoming light from a LED (M530D2, Thorlabs) through the condenser optics, equipped with and without a polarizer for circularly polarized light. Without a polarizer, the illuminating light is unpolarized and the transmitted light is analyzed for linear polarization along 4 linear polarization orientations, i.e. 0°, 45°, 90°, and 135°. With a polarizer in the illumination path, the specimen is illuminated with circularly polarized light, and the transmitted light is again analyzed for the 4 linear polarizations.

The first configuration without a polarizer in the illumination path makes the setup sensitive to diattenuation, while insensitive to the birefringence of the sample. The second configuration with polarizer renders the setup sensitive to both sample properties, diattenuation and birefringence (24). For the study reported here, we used both configurations in order to distinguish between changes in diattenuation and birefringence when analyzing the response of brain tissue slices to stimulation.

In both configurations, the transmitted light with polarizing beam splitters represents the imaging path that projects four images onto the four quadrants of a single EMCCD camera (iXon Ultra, Andor USA, Concord, MA). For the instantaneous PolScope analysis, the four image quadrants undergo the same computational analysis as is done for the polarized fluorescence described in (22). Briefly, for both configurations, the transmitted light intensity recorded in each quadrant is related to the average intensity across the four quadrants (*I*), a polarization factor *p*, and an azimuth orientation *ϕ* as follows:

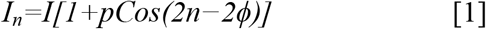

where n=0°,45°,90°,135°.

Retrieval of the average intensity, orientation, and polarization factor of the light through the object is efficiently expressed in terms of the Stokes parameters of the polarization-resolved transmitted light:

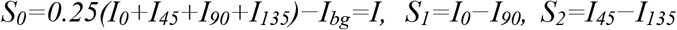

with

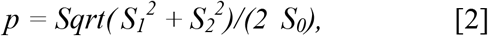

and

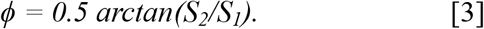

The average intensity I transmitted by the sample is

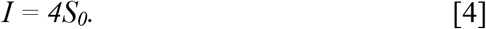

For both configurations, i.e. with and without polarizer in the illumination path, we used the same expressions [2], [3], and [4] for calculating the parameters *p*, *ϕ*, and *I*, but their physical interpretation has to change. For the configuration without polarizer, *p* and *ϕ* are directly related to the diattenuation (*I*_*max*_ – *I*_*min*_)/ (*I*_*max*_ + *I*_*min*_) and the orientation *ϕ* of the linearly polarized light that is maximally transmitted. In the experiments reported in the Results section, we used this configuration to ascertain that the brain tissue has no measurable diattenuation, with or without stimulation. This result was important for the interpretation of the specimen parameters measured with the second configuration setup that included a circular polarizer in the illumination path. If there is no diattenuation in the specimen, then its polarization factor can be interpreted as a measure of linear birefringence in the specimen. In that case, the polarization factor represents a differential retardation measured relative to some fixed retardance.

To quantify the temporal changes in average transmittance and relative birefringence after electrical nerve stimulations, averaged light intensities were calculated at the same selected ROI, which were measured typically at 50 to 70 μm apart from the tip of the stimulating pipette. Peak amplitudes of the signals were evaluated by averaging consecutive 5 frames of single trial, which includes the highest intensity value over the timeframes.

### Statistical analyses

Statistical analyses were performed by using SigmaPlot (version 13.0, Systat, San Jose, CA). All data are expressed as means ± S.E.M. Statistical significance was tested with *t*-test (paired or unpaired), and set significant difference between groups as p<0.05. Normality test was evaluated using the Shapiro-Wilk method. Data that had normality rejected, a Mann-Whitney test was performed as non-parametric test as indicated (p<0.05). Statistical significance among 3 groups was performed by ANOVA, followed by Tukey test when normality test was passed. Data that had normality rejected, a Kruskal-Wallis ANOVA on Ranks was performed, followed by Dunn’s method for pairwise multiple comparison procedures, considered p<0.05 as statistically significant. Data were obtained from at least three different animal preparations.

## Results

### Birefringence map of mouse acute hippocampal slices observed with the LC-PolScope

We observed acute slices of mouse hippocampus by using the Liquid Crystal (LC)-PolScope (Fig. 1). Images of the hippocampal slice, taken with four elliptical polarization illuminations of different principle axes (Figs.1& 2, 0°-S1, 45°-S2, 90°-S3 and 135°-S4) and an image taken with the extinction illumination setting in which the polarization of the illumination has opposite handedness with respect to that of the circular analyzer (Fig. 1B, Fig. 2, S0), were used to obtain the retardance map (Fig. 1C) and the slow axis orientation map (Fig. 1D). The pseudo-colored map of the hippocampus slice in Fig. 1D represents the ensemble slow axis orientation. The averaged image of the four elliptical polarization illumination settings was also shown (Fig. 1A). Intensity profiles of each elliptical polarization illumination setting enclosed by a vertical rectangle in Fig. 1A were plotted in Fig. 1E. From the line profiles of the retardance value, we measured the lowest retardance value of around 3 nm at st. pyramidale, and the highest retardance value of around 10 nm at st. radiatum. The retardance map (Fig. 1C) demonstrated that the dendritic layers of area CA1 (st. radiatum and st. oriens) as the high birefringence area, whereas cell body layers (st. pyramidale and granule cell layer in DG) were observed as the lowest birefringence area. There was a noticeable high birefringence structure at st. lacunosum-moleculare (Fig. 1C, white arrowheads), which represented the axonal projection from the layer III entorhinal cortex neurons to distal dendrites of CA1 pyramidal cells known as temporoammonic system (25). At area CA3, there was a layer with intermediate retardance value compared to the adjacent strata between st. pyramidale and st. radiatum (Figs.1A-D). This area is known as st. lucidum, where many axons from dentate granule cells (mossy fibers) localize along proximal dendrites of CA3 pyramidal cells (26). At st.radiatum of area CA1, slow axis orientation was perpendicular to Schaffer collateral axons and parallel to the main apical dendrites of CA1 pyramidal neurons.

**Figure 1.**
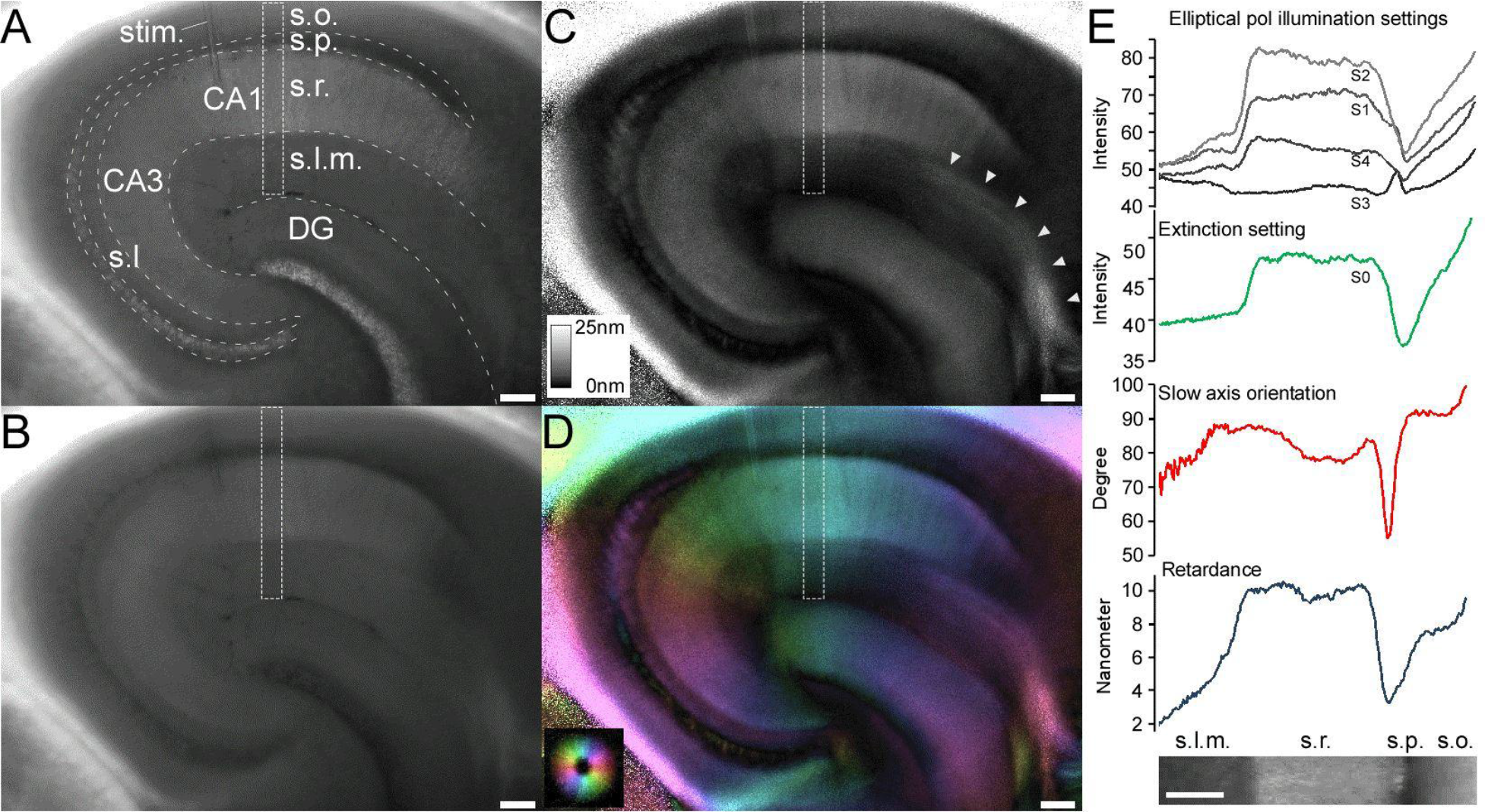
Polarized light imaging of a mouse hippocampal slice using the LC-PolScope. (*A*) An average transmittance image computed from four polarized light images with different elliptical polarization illumination settings. Stratum oriens (*s.o.*), stratum pyramidale (*s.p.*), stratum radiatum (*s.r.*), stratum lacunosum-moleculare (*s.l.m.*), stratum lucidum (*s.l.*). (*B*) A polarized light image with extinction illumination setting (*S0*). (*C*) Retardance map. The local retardance values are proportional to the brightness. The inset shows the reference for local retardance values. (*D*) The slow axis orientation map. The color wheel on the left bottom shows the reference for the local slow axes orientation of birefringence. Bars in *A-D*, 100µm. (*E*) Analysis of optical properties across multiple strata at area CA1 (enclosed by *white broken rectangles* in *A-D*). Intensity profiles are taken with four different elliptical polarization illumination settings (*S1-S4, black*), extinction illumination setting (*S0, green*), slow axis orientation (*red*) and retardance value (*dark blue*). Inset at the bottom is a cropped image from average transmission of area CA1 enclosed by a *white broken rectangle* in *A*. Bar, 100µm.

**Figure 2.**
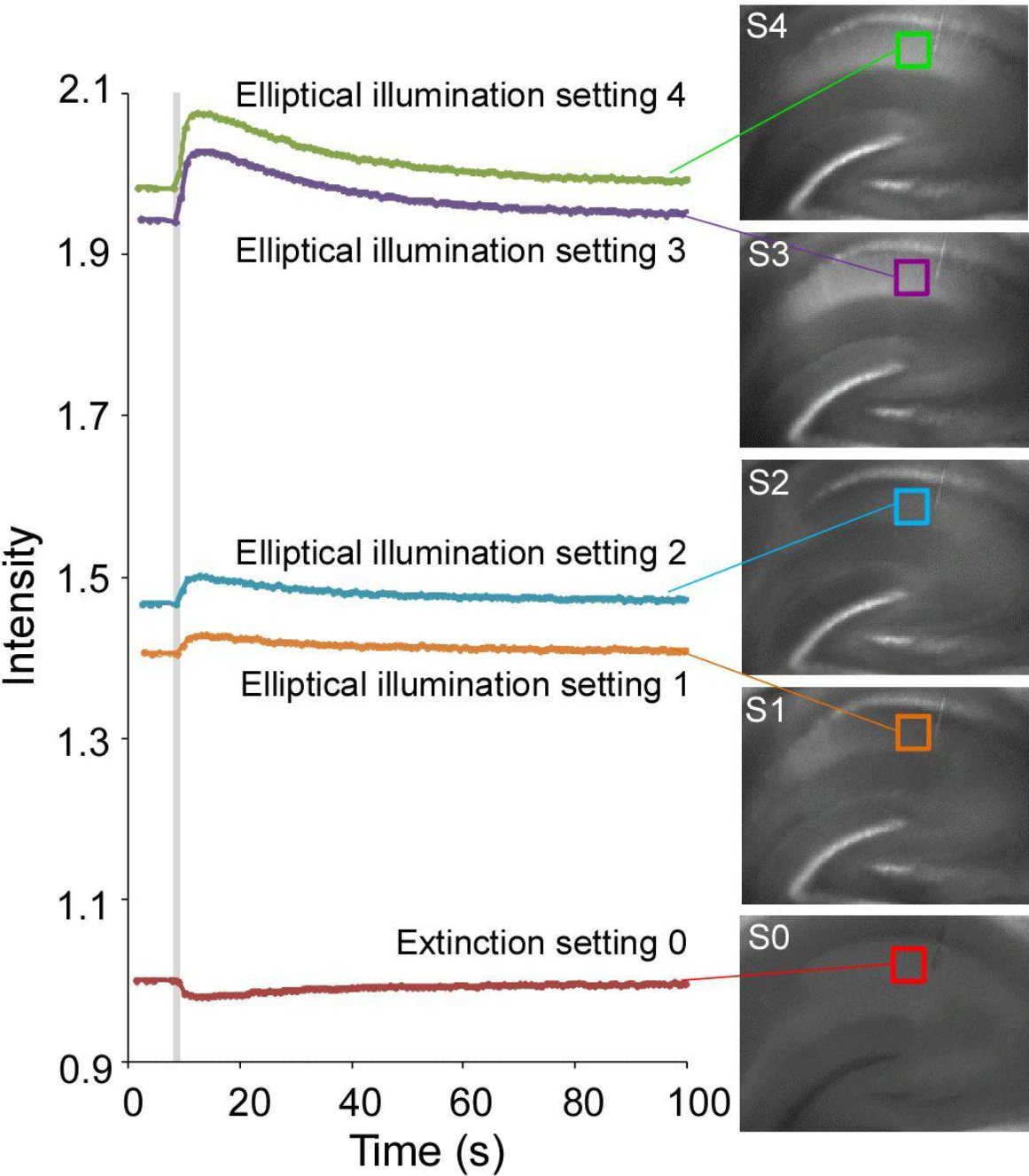
Recording of IOS change using the LC-PolScope measured at stratum radiatum of area CA1 followed by Schaffer collateral stimulation (40 Hz, 50 pulses). The images of a hippocampal slice (*right*) are sequentially acquired with five polarization illumination settings (*setting S0 to S4*). Average intensities of the same area enclosed by colored squares in each image are plotted against the time (*setting 0 in red, setting 1 in orange, setting2 in blue, setting 3 in purple, setting 4 in green*). Electrical stimulation is applied at t=8 sec for 1.25s (shown as a vertical *gray area*). Images of S1 to S4 are obtained with elliptical polarization illumination in LC-PolScope with different principle axes, 0°, 45°, 90° and 135° with respect to the laboratory flame of coordinate. Setting 0 represents extinction illumination condition where the specimen was illuminated with circularly polarized light with the opposite handedness of the circular polarizer used for the analyzer. The intensities are shown as relative values by normalizing the value of setting 0 before the stimulation as 1.0.

### Activity-dependent birefringence changes in hippocampal slices evoked by the electrical stimulation

By using the LC-PolScope, we observed transiently increase of transmittance at st.radiatum of area CA1 by electrical stimulation (40 Hz, 50 pulses) applied at Schaffer collateral axons in all elliptical polarization illumination settings (Fig. 2). In particular, images taken with four elliptical polarization illuminations with principle axes of 0°, 45°, 90° and 135° (Fig. 2, S1, S2, S3, and S4, respectively), a transient *increases* of light intensity in all four elliptical polarization illumination settings were observed after nerve stimulations. These transient increases of transmitted light intensity corresponded to the traditional IOS (5, 7, 9). Interestingly, in the extinction setting, there was a transient *decrease* of light intensity observed after the nerve stimulation (Fig.2, Extinction setting 0 in red). We have not found reasonable explanations for this transient decrease of transmitted light intensity when the slices were observed with extinction illumination setting.

We found that these transmittance changes of polarized light were associated with the birefringence change of the slice. Birefringence change of the hippocampal slice was observed through the sequential acquisition of five images before and after the electrical stimulation over time. Changes in the retardance value at st.radiatum close to the tip of the stimulation electrode (enclosed by red square in Fig. 3A) after nerve stimulation were analyzed as follows (Fig. 3B). At the resting state, the retardance value was 13.31 ± 1.4 nm (n=8). After Schaffer collateral stimulation, the retardance values transiently increased to 13.67 ± 1.4 nm (n=8). The increase of retardance value at the peak was 0.2 – 1.0 % of the resting values (paired t-test, p=0.002). This transient retardance increase was completely diminished in the presence of 1 µM of tetrodotoxin (TTX), a blocker of voltage gated sodium channels (Fig. 3B, lower trace). The stimulation-evoked retardance increase was enhanced in the presence of 40 M of 4-Aminopyridine (4-AP) (Fig. 3C, lower trace), a blocker of slowly inactivating potassium channels (27). These results indicated that the retardance change in st. radiatum is associated with action potentials and following synaptic transmission.

**Figure 3.**
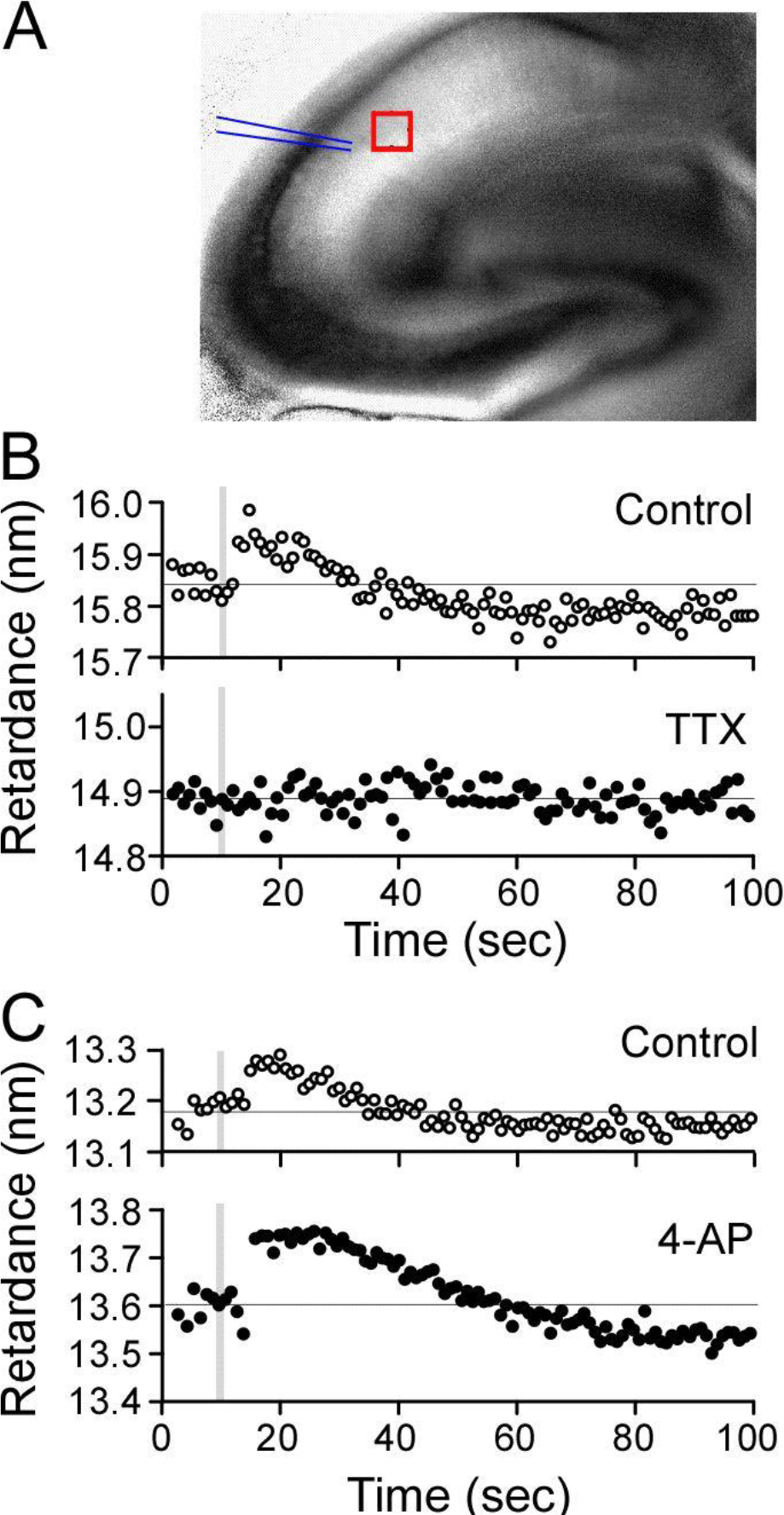
Recordings of retardance change detected by the LC-PolScope. Retardance values are measured at stratum radiatum of area CA1 by Schaffer collateral stimulation (40 Hz, 50 pulses). (*A*) A retardance map of a hippocampal slice. The red square indicates the area for the retardance measurements in B and C. The location of the stimulation electrode is shown as two lines. (*B*) Recordings of the retardance changes in normal ACSF (*Control; open circles*) and in ACSF with 1µM of tetrodotoxin (*TTX; filled circles*) taken at the same location of the hippocampal slice. (*C*) Recordings of the retardance changes in normal ACSF (*Control; open circles*) and ACSF with 40µM of 4-aminopyridine (*4-AP; filled circle*) from the same hippocampal slice. The timings of electrical stimulations (40 Hz, 50 pulses) are shown as vertical gray areas in the graphs. Thin solid lines in the graphs indicate the resting retardance value obtained by an average of 10 frames at the resting level before the stimulation.

### Birefringence map of single neurons in primary cultures

The retardance map of hippocampal slices was derived from the ensemble value in the optical path length of transmitted light through the thickness of slice preparations (thickness, 250 µm). To obtain the retardance value and the slow axis orientation of single neurons, we analyzed the polarized light images of principal neurons in primary neuronal cultures. In these observations, we used 60x 1.4NA oil immersion lens (Nikon PlanApo VC) that allows sub-cellular birefringence mapping of the principal neurons (Fig. 4). Slow axis orientation map of single neurons revealed that there were two different birefringent components at the dendritic structure, which were observed at the core and near the surface of the dendritic shafts. The birefringent structure that filled inside the dendrites (Fig. 4B, magenta) had slow axis orientations parallel to the longitudinal axis of the dendrite. The retardance value of the cytoplasmic structures inside the dendrite was 1.04 ± 0.10 nm (n=8). The cytoplasmic birefringent component was surrounded by another birefringent component along the surface of the dendrites (Fig. 4B, blue-green). This sheathe-like birefringent component had slow axis orientation perpendicular to the length of the dendrite. This birefringence component had retardance values of 0.51 ± 0.03 nm (n=8). These observations were consistent with the previous observations of birefringent structures in the neuronal protrusions of *Aplysia* bag cell neurons using the LC-PolScope (20, 28).

**Figure 4.**
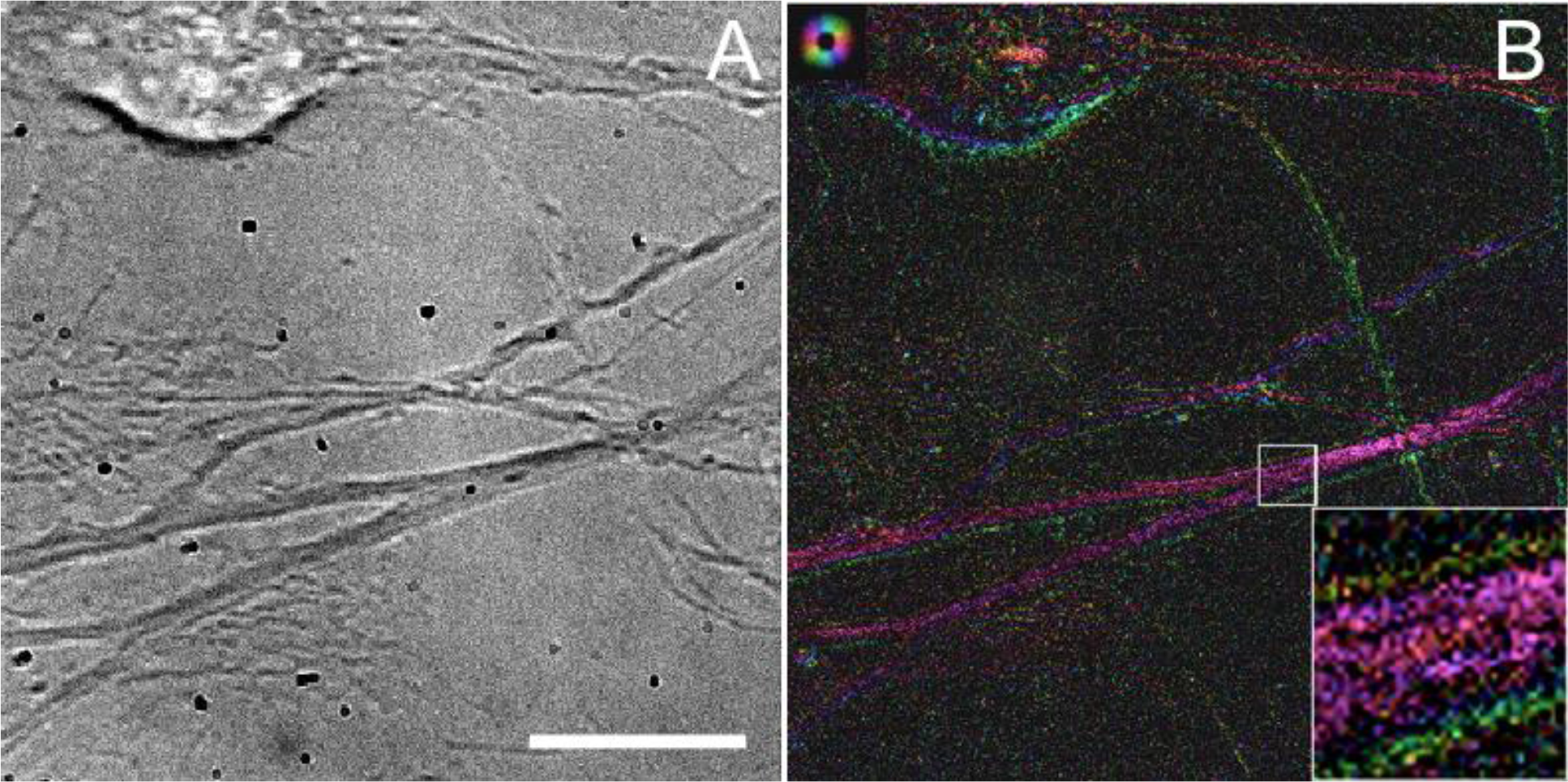
The LC-PolScope image of primary cultured hippocampal neurons. Transmitted light image (*A*) and the slow axis orientation map (*B*) of a cell body (top left) and dendrites (middle) are shown. An area enclosed by a white square in *B* is magnified as an inset at the right-bottom corner. The color wheel on the top left shows the reference for the slow axis orientation. Bar, 10µm.

### Instantaneous PolScope imaging to study the mechanisms of activity-dependent birefringence changes in mouse hippocampal slices

In our LC-PolScope imaging, constant transmittance through the specimens is a prerequisite for the computational analysis of birefringence. Since the LC-PolScope sequentially acquires five images of different polarization illumination settings, rapidly changing transmittance causes unavoidable artifact in calculating the retardance value and the slow axis orientation. To avoid this artifact and to achieve high temporal resolution to examine the activity-dependent birefringence changes of mouse hippocampal slices, we developed the instantaneous PolScope, which was the modified version of our instantaneous FluoPolScope, that had been developed for tracking position and orientation of single fluorophores in living cells (22). Hippocampal slices were illuminated with circularly polarized light, and the instantaneous polarization analysis was done by re-collimating the light originating from the primary image plane of the microscope. In the collimated space, the light is first divided equally between two arms by a polarization independent beam splitter (50:50 mirror in Fig. 5A, 21014, Chroma Technology). A pair of polarization beam splitters (PBS, Fig.5A, Semrock) separated the light into four images and analyzes their linear polarization along 0°, 45°, 90° and 135° orientations. One PBS generates 0° and 90° polarization beam paths, while the polarization in the other arm was first rotated by 45° using a half-wave plate (HWP in Fig.5A, Meadowlark Optics NH-532) and then passed through the other PBS to generate the 45° and 135° paths. Subsequently, we use broadband mirror-based beam-steering optics and a focusing lens to project four images onto the four quadrants of a single EMCCD detector (Fig. 5B). We detected polarized light intensity changes through the hippocampal slices in all four illumination states (I_0°_, I_45°_, I_90°_, I_135°_), when Schaffer collaterals were stimulated (Fig. 5C). From the acquired images we computed the polarization factor *p*, hereafter this value will be referred as “relative birefringence” (Fig. 5D, top), and the average transmittance of all quadrant images (Fig. 5D, bottom) before and after the neuronal activation over time. Thus, by using the instantaneous PolScope imaging, we could observe the temporal change of relative birefringence map that corresponded to the retardance map obtained by the LC-PolScope, with high time resolution. The time resolution of this system was determined by the acquisition rate of the camera, and was approximately 100Hz for our EM CCD camera (Andor iXon Ultra). Compared to the LC-PolScope, the instantaneous PolScope improved the time resolution by a factor of 100. With these improvements, polarized light microscopy is ready for prime time in analyzing the intrinsic polarization signals of neurons.

**Figure 5.**
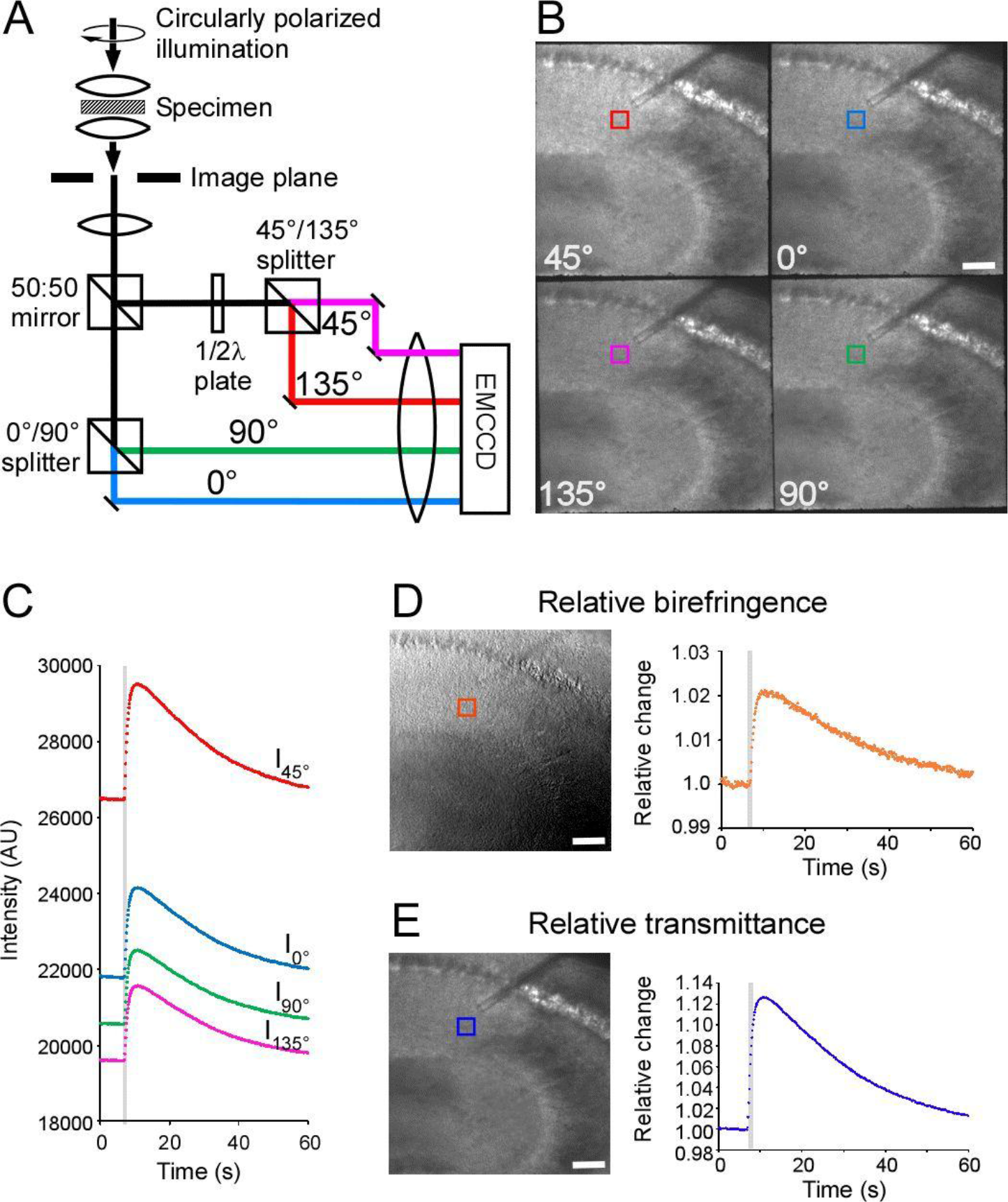
The instantaneous PolScope. (*A*) A schematic drawing of the four-way linear polarization splitting system for the instantaneous analysis of polarized light through the specimen. (*B*) Quadrant images of mouse hippocampal slice captured by an EM CCD camera. Orientations (bottom left) of each image indicate the transmission axis of the polarization analysis. Bar, 100µm. (*C*) Light intensity changes of a selected area in each quadrant image at stratum radiatum of area CA1 (enclosed by squares in *B*) are plotted against time. The timing of electrical stimulation (40Hz, 50 pulses) was indicated by a gray vertical line. (*D, left*) Relative birefringence image computed from four images shown in *B*. (*D, right*) Change in the relative birefringence at the selected area (*orange square* shown in *D*). (*E, left*) Average transmittance image from four images shown in *B*. (*E, right*) Change in the average transmittance at the selected area (*purple square* shown in *D*). Bars, 100 µm.

### Birefringence versus diattenuation of hippocampal slices observed with the instantaneous PolScope

The polarization-sensitive IOS of hippocampal slices could be derived either from the birefringence change or the diattenuation change, or the combination of both. We examined the contribution of diattenuation, a polarization-dependent absorption, to the polarization-sensitive IOS in response to nerve stimulations (24). To test these two polarization sensitive optical properties, we illuminated the hippocampal slice with two different illumination conditions; 1) circularly polarized illumination (Fig. 6A) and 2) non-polarization illumination (Fig. 6B). Ideally, the diattenuation change can be detected in both illumination conditions, whereas the birefringence change can be detected only with the circular polarization illumination condition. The stimulation-evoked polarization-sensitive IOS (pol-IOS) change was detected when the brain slice was illuminated with circularly polarized light (Fig. 6C), but not with non-polarized illumination (Fig. 6D). This result clearly indicated that the pol-IOS change at area CA1 was originated predominantly from transient changes of birefringence, not from the changes in diattenuation.

**Figure 6.**
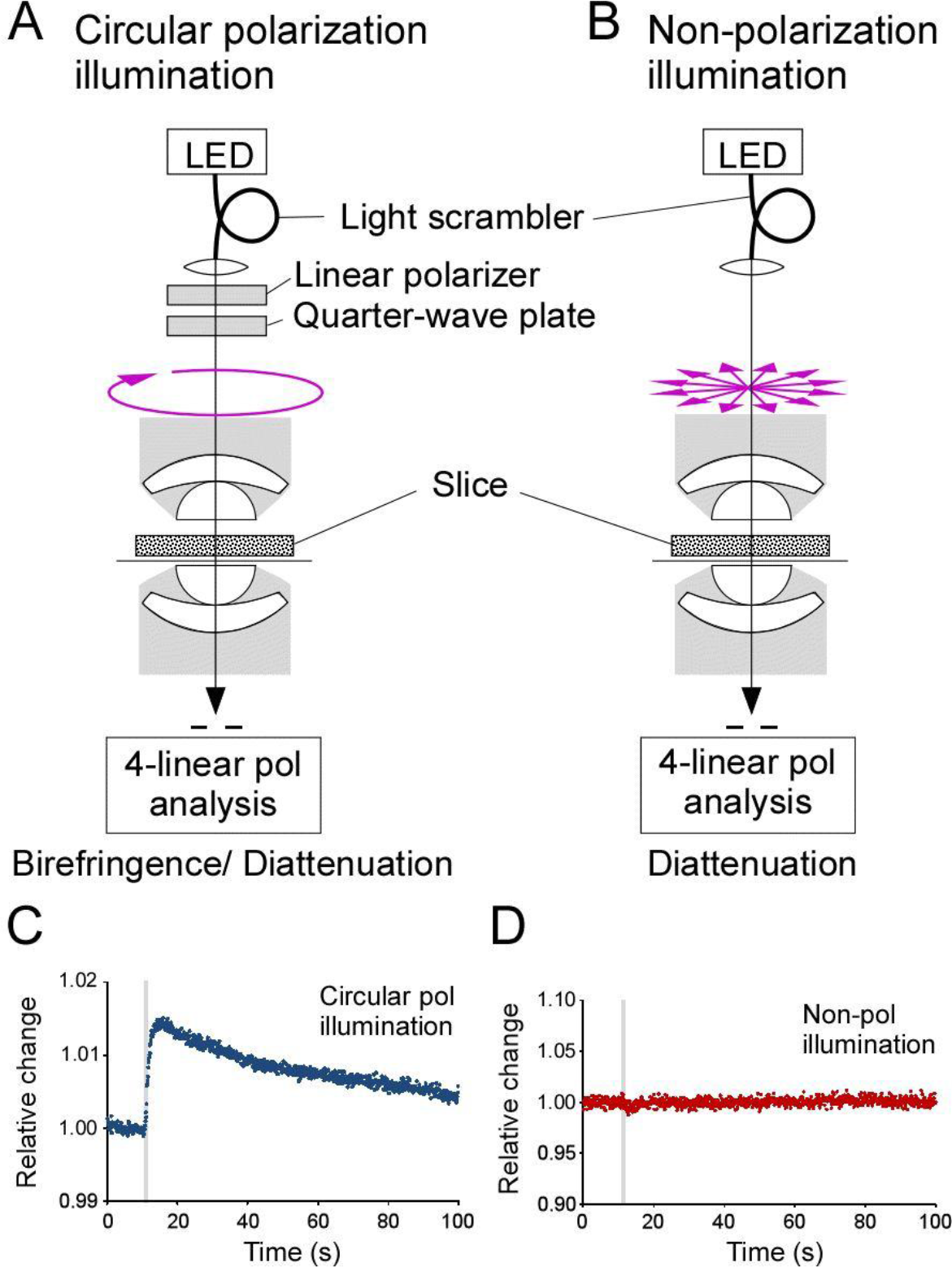
Birefringence versus diattenuation as origins of polarization sensitive IOS. (*A*) A schematic of optical system with circular polarization illumination that detects both birefringence and diattenuation changes of the specimens. (*B*) A schematic of optical system with non-polarizing illumination that detects diattenuation change but does not detect birefringence change of the specimens. (*C, D*) Changes in the relative birefringence (*C*) and diattenuation (*D*) of the same hippocampal slice before and after the electrical stimulations (*gray vertical lines*).

### Spreading of stimulation-evoked birefringence changes at CA1 of hippocampal slices

The spatial propagation of relative birefringence change was observed at st.radiatum of area CA1 after Schaffer collateral stimulation. To observe larger field of view, we used a low magnification objective lens (XL Fluor 4x, NA0.28) (Figs. 7A&C). To enhance the changes, intensities of both transmittance (IOS) and birefringence (pol-IOS) images at each frame were subtracted by the images of their resting state before the stimulation. Following the electrical stimulations, the transmittance change spread along st. radiatum (Fig. 7B). This result was consistent with the IOS imaging of hippocampal slices reported by others (5, 9, 13, 29). Similarly to the transmittance changes, we observed spreading of birefringence initiated from the site of electrical stimulation (Fig. 7D). The birefringence signal spread along st. radiatum toward both CA3 and subiculum directions, but did not spread to other strata, such as st.oriens and st. lacunosum-moleculare. Interestingly, around 60 sec after the stimulation when birefringence signal diminished at st. radiatum, a weak birefringence increase was observed at st. pyramidale (Fig. 7D), where the cell bodies of CA1pyramidal neurons were located.

**Figure 7.**
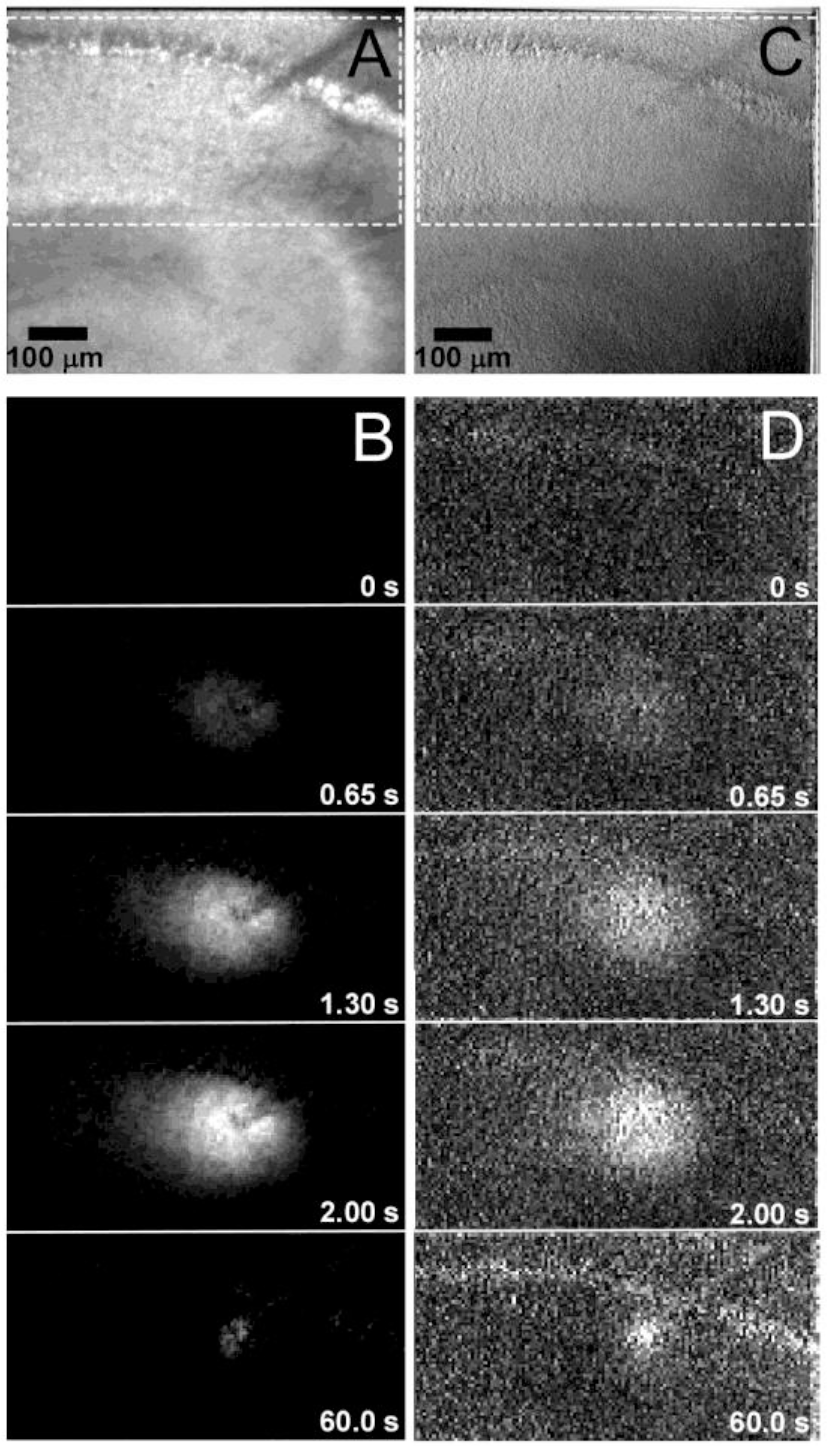
Spreading of traditional IOS and polarization-sensitive IOS at stratum radiatum of area CA1 taken by the instantaneous PolScope. (*A*) An average transmittance image of the hippocampal slice. A glass pipette for electrical stimulation is seen at the *right top<* of the image. (*B*) Spreading of the traditional IOS in transmitted light imaging observed in the area enclosed by broken white rectangle in A. Numbers shown at the *right bottom* show the time after the electrical stimulation applied at t=0s (40Hz, 50 pulses). (*C*) A relative birefringence image of the hippocampal slice. (*D*) Spreading of polarization sensitive IOS change observed in the area enclosed by broken white rectangle in C. To enhance the stimulation evoked changes of the images, in B and D, the averaged images before the stimulation were subtracted from all the frames. Bars, 100 µm.

### Local difference of stimulation-evoked birefringence changes at stratum radiatum of area CA1

Through the correlative observations of pol-IOS and traditional IOS, we explored the sub-cellular structures at st.radiatum of area CA1 where the activity-induced birefringence changes were produced. Through the instantaneous PolScope analysis, the time series of transmittance map (Fig. 8A) and that of the relative birefringence (Fig. 8B) before and after the Schaffer collateral stimulation were obtained. Based on the value of birefringence at the resting state, two segmented zones were created referring high resting birefringence (green areas in Fig. 8C) and low resting birefringence (blue areas in Fig. 8C). We found local difference of stimulation-evoked birefringence changes (Fig. 8E), which correlated with the local difference of sub-cellular birefringence. We observed larger pol-IOS predominantly at locations where the resting birefringence values were higher than the surrounding locations (Fig. 8C). Interestingly, the local difference that we detected with pol-IOS was not clearly observed with the traditional IOS imaging (Fig. 8D). These observations indicated that the local difference of pol-IOS derived from the difference of static birefringent structures in the slices.

**Figure 8.**
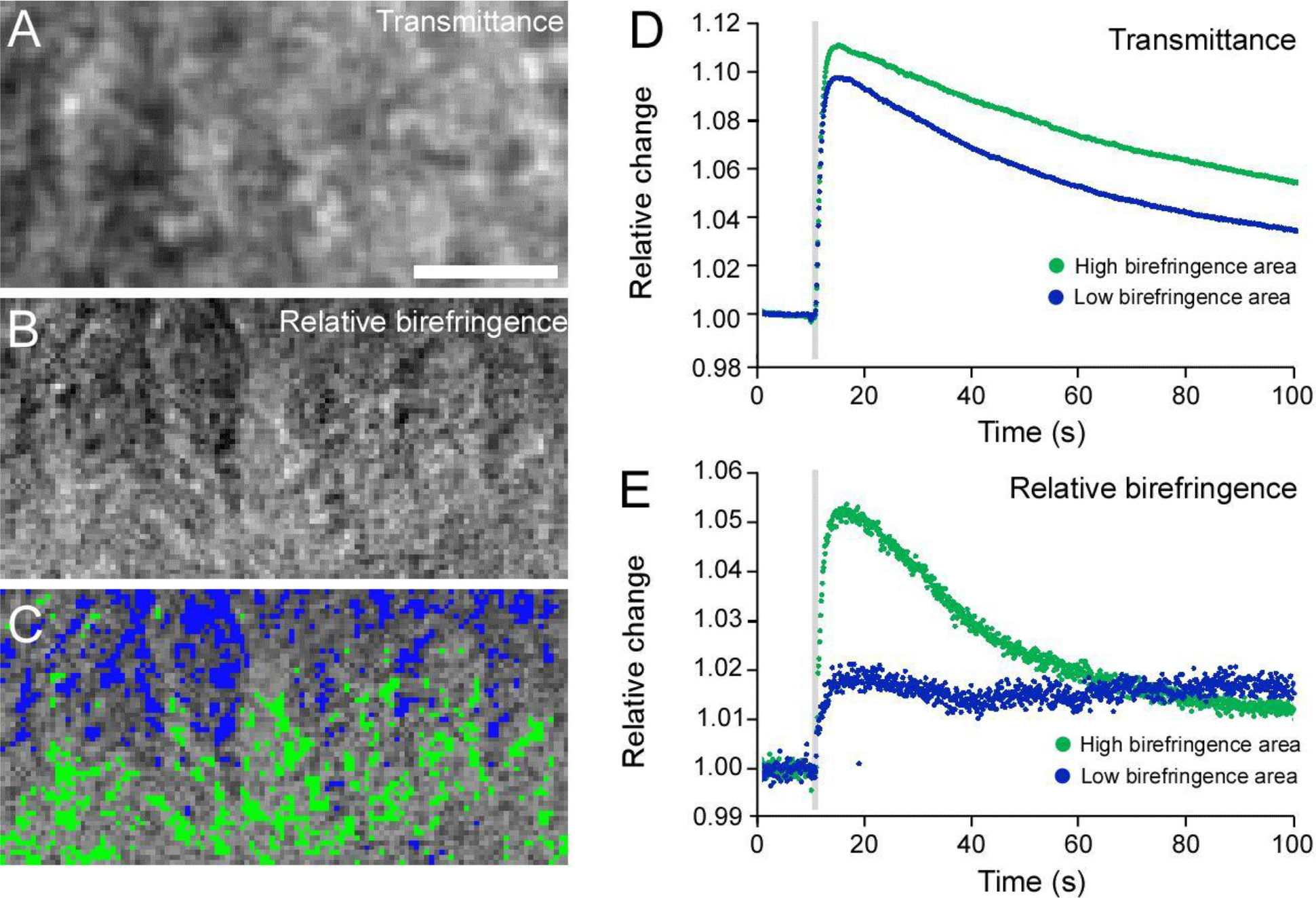
Local difference of polarization sensitive IOS at stratum radiatum of area CA1. (*A*) Average transmittance map. Bar, 10µm. (*B*) Relative birefringence map. (*C*) Segmentation of high birefringence area exhibiting top 10 % (Green) and low birefringence area exhibiting bottom 10 % (Blue) of the brightness distribution at areas shown in *B*. (*D, E*) The average transmittance change (*D*) and relative birefringence change (*E*) observed at high birefringence regions (green) and low birefringence regions (blue) are plotted against time, respectively. Electrical stimulation (40Hz, 50 pulses) is applied during the time shown in gray vertical areas. Values (*Relative change*) in *D* and *E* are normalized so that the average values before the simulations are 1.0.

### Correlation between the number of stimulation pulses and the amplitudes of pol IOS or traditional IOS

To characterize the temporal changes of pol-IOS by different number of stimulation pulses, various numbers of current pulses (amplitude, 250µA; duration, 1ms) were applied at 40 Hz to Schaffer collateral. We measured changes in the relative birefringence and the transmittance in a selected square area (approximately 800 µm^2^) in st. radiatum before and after the stimulations (Fig. 9). A small, but detectable relative birefringence change was observed by a single pulse stimulus. The relative birefringence change increased as the number of stimulation pulses increased, and reached nearly saturating level at 50 pulses (Fig. 9A, B). The stimulation-response curve of relative birefringence change was similar to that of the transmittance change (Fig. 9 C,D). Based on these results, we used 50 pulses as standard electrical stimulation hereafter to induce excitation of Schaffer collaterals and following excitatory synaptic transmission between CA1 pyramidal neurons. The evoked birefringence and transmittance signals were well reproducible in peak amplitude, rise time, and decay time.

**Figure 9.**
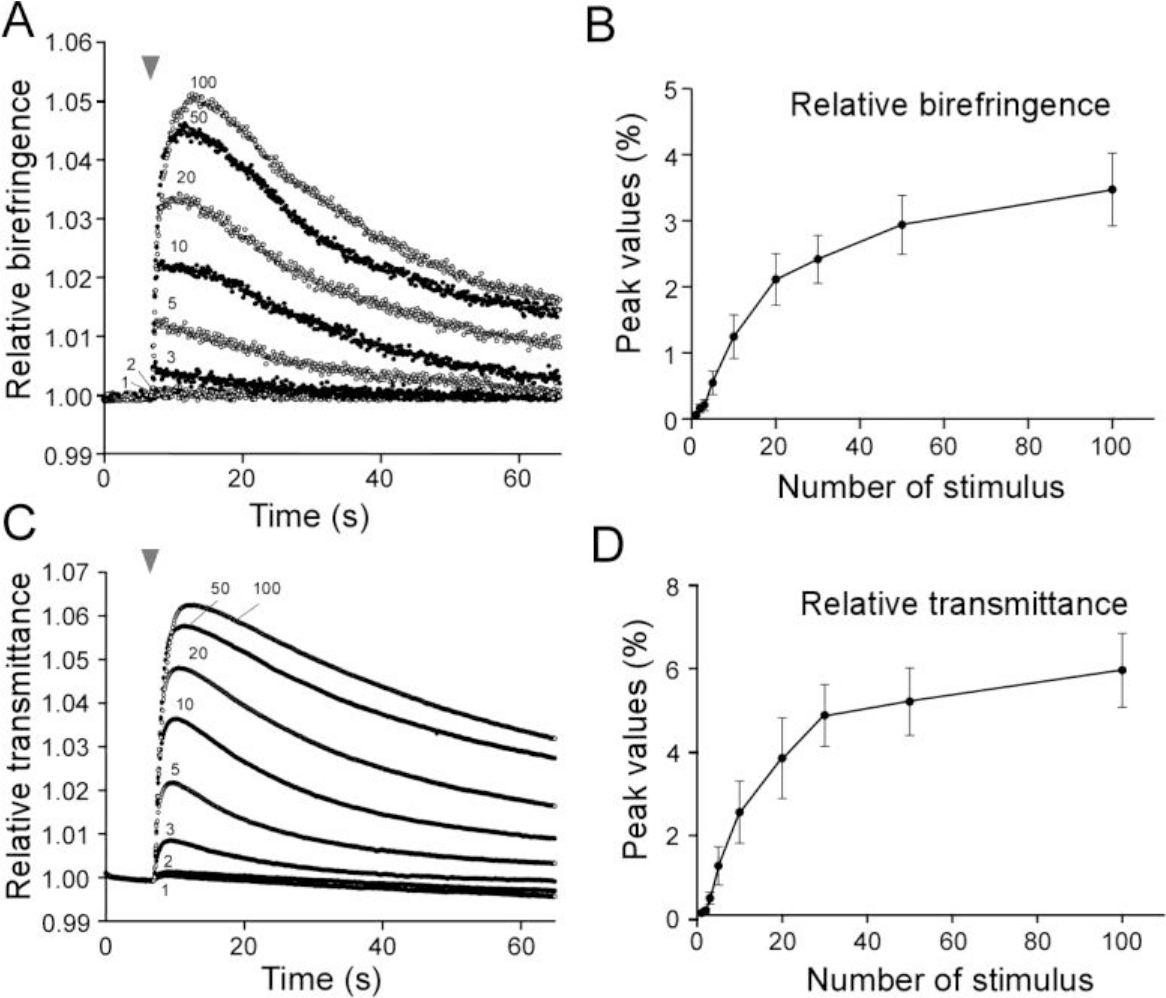
Relationship between the number of stimulation pulses and the IOSs changes. (*A*) Superimposed traces of relative birefringence changes elicited by various numbers of pulses (*1, 2, 3, 5, 10, 15, 20, 50* and *100*) with 1.25ms intervals (40Hz). (*B*) Peak values of relative birefringence were plotted against the applied number of pulses. *Arrowhead* indicates the start of stimulation. (*C*) Superimposed traces of relative transmittance changes elicited by various numbers of stimulation pulses that correspond to the relative birefringence changes in *B*. (*D*) Peak values of relative transmittance were plotted against the applied number of pulses. Dara are shown in mean ± S.E.M. (n=6).

### Distinct decay time courses of temporal changes in birefringence and transmittance signal

We have quantitatively compared the temporal changes of relative birefringence and transmittance of the same area of stratum radiatum of area CA1 induced by electrical stimulation. Fig. 10 shows representative relative birefringence and average transmittance signals obtained from the same datasets followed by train of 50 pulses of stimulation (1ms duration) at 40Hz. Rise time (20 % − 80 %) of birefringence and average transmittance were 1.44 ± 0.13 sec and 1.21 ± 0.064 sec respectively, indicating no significant difference in rise time between these two IOSs (Mann-Whitney test, p=0.77, n=30). Decay time of relative birefringence and average transmittance were evaluated by the value of time constant obtained by single exponential function fitted between 80 – 20 % of the peak amplitude (solid red lines on each sample traces, τ = 29.6 sec for birefringence, and τ = 48.3 sec Fig. 10A &B). There was a slight variation in the peak amplitude of birefringence among brain slices, probably due to the slight variation of the thickness and the angle of the tissue, or caused by other factors that affects net light intensity through optical light path. However, no correlation was found between peak amplitude of birefringence and its decay time constant obtained by each individual trace (data not shown). For most of the pairs (25 out of 28 trails) of transmittance and birefringence data sets, the decay time constant of birefringence was faster than that of transmittance (Fig. 10C). The paired plots of decay time constants of birefringence and transmittance signals were fitted by a regression line with a slope of 1.83. Averaged value of time constant was 30.3 ± 2.3 sec for birefringence and 55.5 ± 7.1 sec for transmittance, indicating significant difference in the speed of decay time (Fig. 10D) (Mann-Whitney test, p<0.001, n=28).

**Figure 10.**
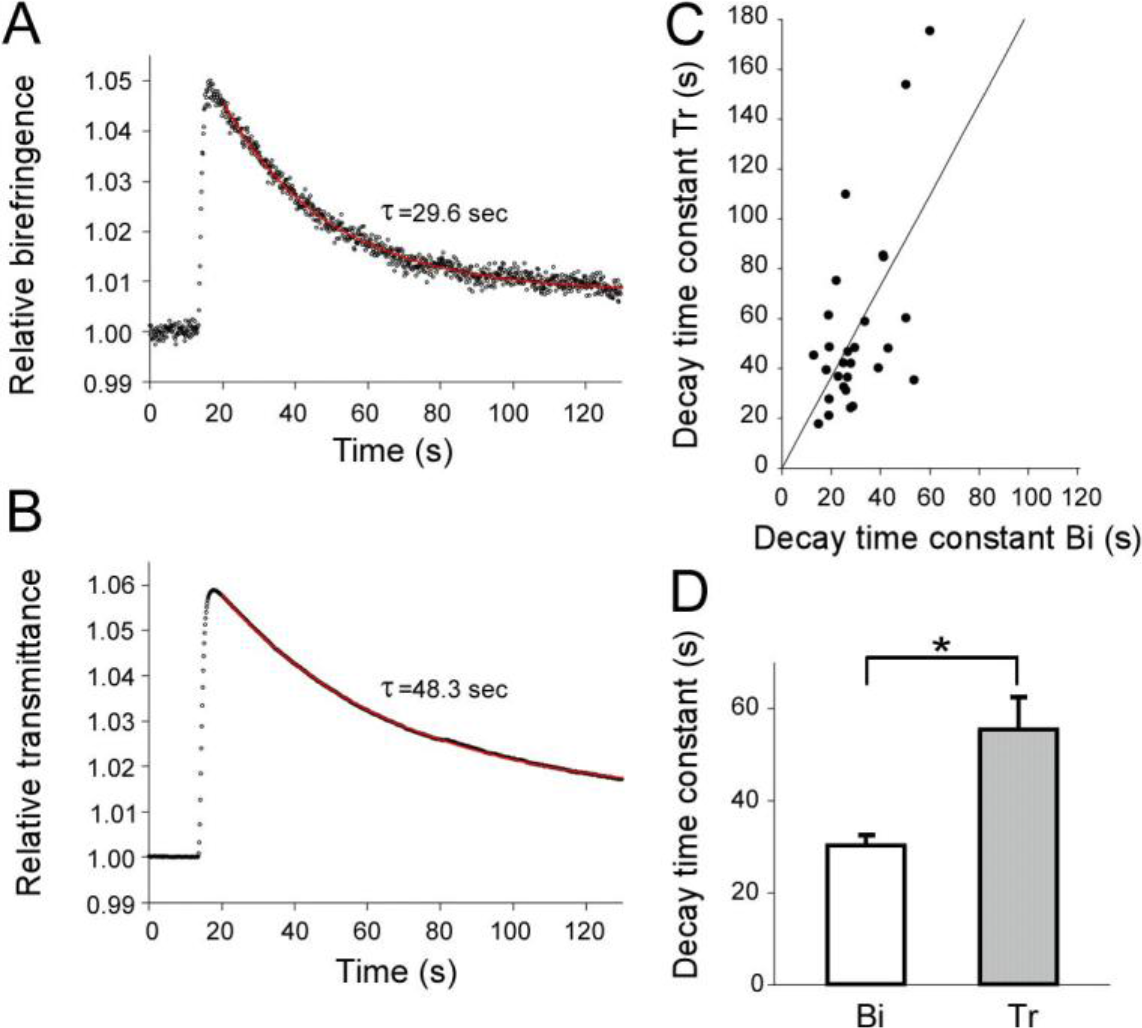
Decay time analysis of IOSs. (*A*) Relative birefringence change measured at stratum radiatum of area CA1 (*black dots*). Decay time constant was estimated by fitting the decay phase of relative birefringence (80 − 20 % of amplitude) with single exponential function (*red solid line*). (*B*) Relative transmittance change of the same specimen and area in *A*. (*C*) Decay time constants of the polarization sensitive IOS (r*elative birefringence; Bi*) and the traditional IOS (*average transmittance; Tr*) obtained from the same response are plotted as a pair. There is a positive correlation between *decay time constant of Bi* and *Tr* (the slope of the regression line is 1.83, r=0.6, p<0.001, n=28, linear regression analysis), (*D*) Averaged decay time constant of relative birefringence (Bi) and average transmittance (Tr). The time constants for relative birefringence was 30.3 ± 2.35 sec (*open bar*) and 55.5 ± 7.09 sec for relative transmittance (*gray bar*). Data are shown in mean ± SEM. Significant difference is found between Bi and Tr (* Mann-Whitney test, p<0.001, n=28).

### Glutamate as a key molecule for generating activity-dependent birefringence changes in hippocampal slices

Larger birefringence changes were typically observed at st. radiatum, st. orience and st. lacunosum-moleculare in area CA1 followed by electrical stimulations. These strata include many synaptic inputs from axons that make synapses between apical and basal dendrites of CA1 pyramidal neurons. In particular, st. radiatum is known to have dendritic alignments lining radially from st. pyramidale (30), receiving many excitatory synaptic inputs that transmit glutamate as a neurotransmitter from axons of CA3 pyramidal neurons.

To address whether an external glutamate application alone can elicit birefringence change, 10 mM glutamate was focally applied to st. radiatum through a glass micropipette (Fig. 11). A transient increase of birefringence was observed after puff application of glutamate for 1s (Fig. 11). This glutamate-evoked transient increase in birefringent was not affected by adding 1 μM TTX in ACSF, indicating that postsynaptic activity alone can generate transient birefringence changes at st. radiatum in the hippocampal slice.

**Figure 11.**
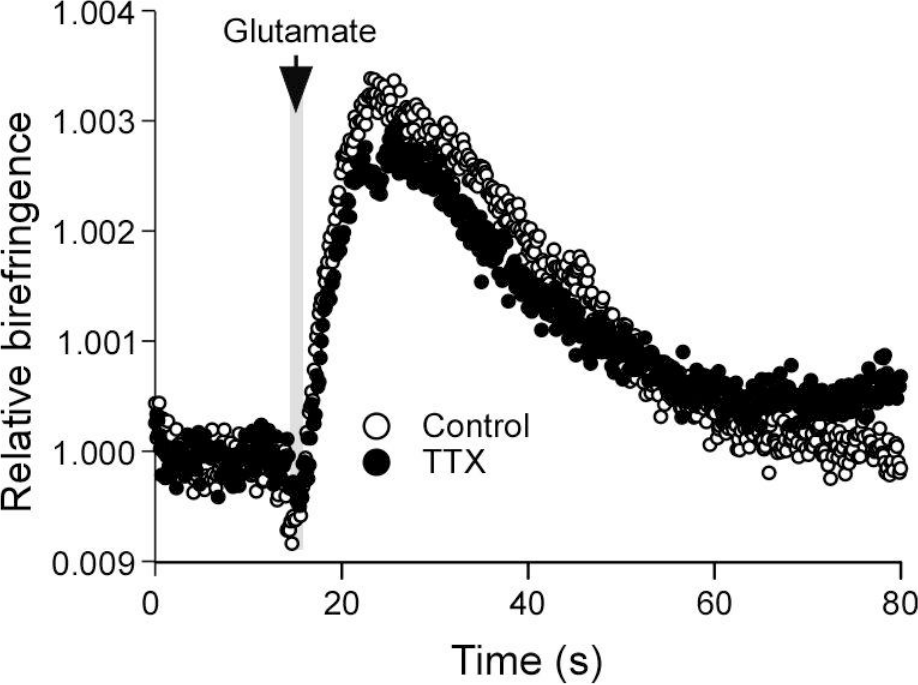
Generation of Pol-IOS by external glutamate application to hippocampal slice both in the presence and absence of TTX. A pulse application of 10mM glutamate through a glass pipette for 1 sec duration (indicated by *gray area*) caused relative birefringence changes both in the presence (*control; open circle*) and absence of 1µM Tetrodotoxin (*TTX; filled circle*).

To further address the mechanisms underlying generation of stimulation-evoked birefringence changes in the hippocampal slice, involvement of excitatory synaptic transmission was examined by pharmacological approaches. First, involvement of ionotropic glutamate receptors on the generation of birefringence and transmittance changes was tested. Both AMPA-type and NMDA-type glutamate receptors are abundantly expressed at spines and dendrites of pyramidal neurons, which are known to play major role on excitatory synaptic transmission and plasticity in hippocampus (31, 32). 10 µM of 6-cyano-7-nitroquinoxaline-2,3-dione (CNQX), a potent and selective blocker of AMPA-type glutamate receptors, decreased the amplitude of relative birefringence of 3.98 ± 0.57 in control to 2.65 ± 0.34 in CNQX (Fig. 12A, B, amplitudes are shown as % of the resting values). 50 μM of (*2R*)-amino-5-phosphonovaleric acid (D-APV), a selective blocker of NMDA-type glutamate receptor, in addition to 10 µM of CNQX, induced significant suppression of the amplitude of relative birefringence changes (CNQX+APV, 1.28 ± 0.18, ANOVA p<0.001, n=7, Fig. 12B). The decay time constant became slightly faster from 29.6 ± 3.90 s in control to 25.3 ± 2.77 s in CNQX, and significantly became faster to 14.2 ± 1.71 s with additional application of D-APV (ANOVA, p=0.018, n=7, Fig. 12C). The amplitude of the relative average transmittance decreased from 6.69 ± 0.78 (control) to 5.23 ± 0.48 in ACSF with 10 μM CNQX, and was significantly decreased to 1.92 ± 0.38 by addition of 50 μM D-APV to ACSF with 10 μM CNQX (ANOVA, p=0.003, Fig. 12E, amplitudes are shown as % of the resting values). The decay time constant of this average transmittance became slightly faster from 69.6 ± 15.8 s (control) to 51.1 ± 8.45 s (CNQX), and further accelerated to 27.6 ± 3.30 s in the presence of D-APV, which was significant change (Kruskl-Wallis ANOVA rank, p=0.005, n=7, Fig. 12F). These data indicated that 70 % and 72 % of birefringence and transmittance signals, respectively, were suppressed by the inhibitors for AMPA and NMDA receptors. Our results indicated that birefringence and transmittance signals induced by electrical stimulation were generated predominantly through the mechanisms that involved these postsynaptic glutamate receptors. The decay time was accelerated significantly in the presence of D-APV (48 % and 40 % of control in polarization anisotropy and transmittance, respectively) but not CNQX alone, suggesting that NMDA receptor and AMPA receptor are differently involved in the generation of IOSs.

**Figure 12.**
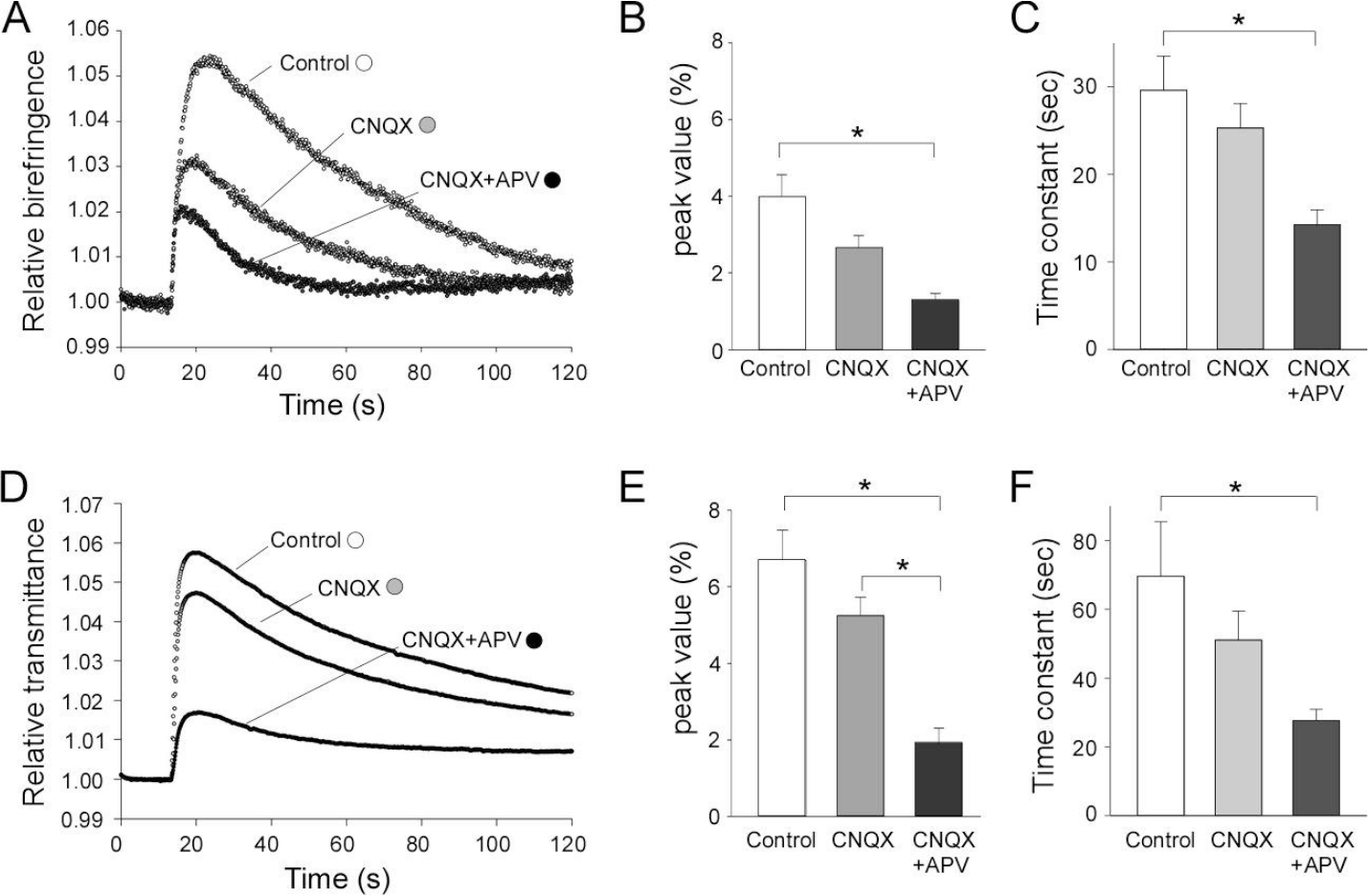
Effects of ionotropic glutamate receptor antagonists on activity dependent relative birefringence change and transmittance change at stratum radiatum of area CA1. (*A*) A series of recordings of relative birefringence changes induced by electrical stimulation in ACSF (*control; open circle*), with 10 µM CNQX (*CNQX; gray circle)*, and with 10 µM CNQX and 50 µM D-APV (*CNQX + APV; filled circle*). (*B*) Summary of the effects of CNQX and D-APV on the peak amplitude of relative birefringence change. The peak value is expressed as increase in % from the resting birefringence value. 3.98 ± 0.57 % (*control; white bar)*, 2.65 ± 0.34 % (*CNQX; gray bar*) and 1.28 ± 0.18 % (*CNQX+APV; black bar*). Significant change is found between control and CNQX+APV (p<0.001, n=7), one-way ANOVA followed by pairwise multiple comparison procedures (*post hoc* Tukey test). (*C*) Summary of the effects of CNQX and D-APV on decay time constant of relative birefringence change. Decay time constant is estimated by fitting single exponential function. 29.6 ± 3.90 sec (*control; white bar*), 25.3 ± 2.77 sec (*CNQX; gray bar*) and 14.2 ± 1.71 sec (*CNQX+APV; black bar*). Significant change is found between control and CNQX+APV (p=0.018, n=7), one-way ANOVA followed by pairwise multiple comparison (*post hoc* Tukey test). (*D*) A series of recordings of relative transmittance changes induced by electrical stimulation in normal ACSF (c*ontrol; open circle*), with 10 µM CNQX (*CNQX; gray circle*), and with 10 µM CNQX and 50 µM D-APV (*CNQX + APV; filled circle)*. (*E*) Summary of the effects of CNQX and D-APV on the peak amplitude of relative transmittance change. The peak value is expressed as increase in % from the resting transmittance value. 6.69 ± 0.78 % (c*ontrol; white bar*), 5.23 ± 0.48 % (*CNQX; gray bar*) and 1.92 ± 0.38 % (*CNQX+APV; black bar*). Significant changes are found between control and CNQX+APV (p<0.001, n=7), CNQX and CNQX+APV (p=0.003, n=7), one-way ANOVA followed by pairwise multiple comparison procedures (*post hoc* Tukey test). (*F*) Summary of the effects of CNQX and D-APV on the decay time constant of relative transmittance change. 69.6 ± 15.8 sec (*control; white bar*), 51.1 ± 8.45 sec (*CNQX; gray bar*) and 27.6 ± 3.30 sec (*CNQX+APV; black bar*). Significant change is found between control and CNQX+APV (p=0.005, n=7), Kruskl-Wallis one-way ANOVA on Ranks followed by pairwise multiple comparison procedures (Dunn’s method). Data are expressed as mean ± S.E.M. *p<0.050.

To further elucidate the processes and mechanisms underlying the generation of birefringence in the hippocampal slice, we tested the effects of blockers of glutamate transporters in neurons and glial cells. Rapid clearance of glutamate from the synaptic cleft is critical for preventing excitotoxicity and for maintaining reliability of excitatory synaptic transmission. Glutamate uptake is mediated by glutamate transporters located at plasma membranes of neurons and astrocytes (33). There are three types of glutamate transporters, GLAST, GLT-1 and EAAC1 (rodent analogs of the human transporters EAAT1, 2 and 3, respectively) that are known to be present in the mature hippocampus. GLT-1 and GLAST are expressed predominantly in plasma membranes of astrocytes. GLT-1 is known to play the major role for clearance of extracellular glutamate near excitatory synapses (34). It is reported that GLT-1 is also detected in axon terminals of CA3 pyramidal cells in the hippocampus, and take up extracellular glutamate into nerve terminals (35, 36). EAAC1 is a neuron-specific glutamate transporter, which is abundantly expressed at postsynaptic density (34, 37), dendritic shafts and spines surrounding active zones of excitatory synapses in CA1 pyramidal neurons (38). We used two different inhibitors of glutamate transporters to address the contribution of glial and neuronal glutamate transporters to the generation of birefringence change accompanied by neuronal activation. Dihydrokainate (DHK) is a selective blocker of glial glutamate transporter GLT-1 (39). DL-*threo*-β-Benzyloxyaspartic acid (TBOA) blocks both neuronal and glial glutamate transporters at lower concentration on GLT-1 and EAAC1 than glial transporter, GLAST (40). To compare the involvement of GLT-1 and EAAC1 in birefringence and transmittance signal, we applied 50 μM of DL-TBOA, or 300 μM of DHK to hippocampal slices, and compared before and after the application of those blockers. Relative birefringence was dramatically enhanced by DL-TBOA (Fig. 13A), whereas DHK had little effect (Fig. 13B). The average increase of peak amplitude (in %) from the resting values analyzed from pooled data demonstrated birefringent signal increased significantly from 6.74 ± 2.10 (control) to 13.12 ± 3.76 (DL-TBOA) (Fig, 13C, paired *t*-test, p=0.036, n=5). In contrast, DHK induced no significant change (Fig. 10D, 8.42 ± 2.66 in control, 7.88 ± 2.54 in DHK, paired *t*-test, p=0.36, n=5).

**Figure 13.**
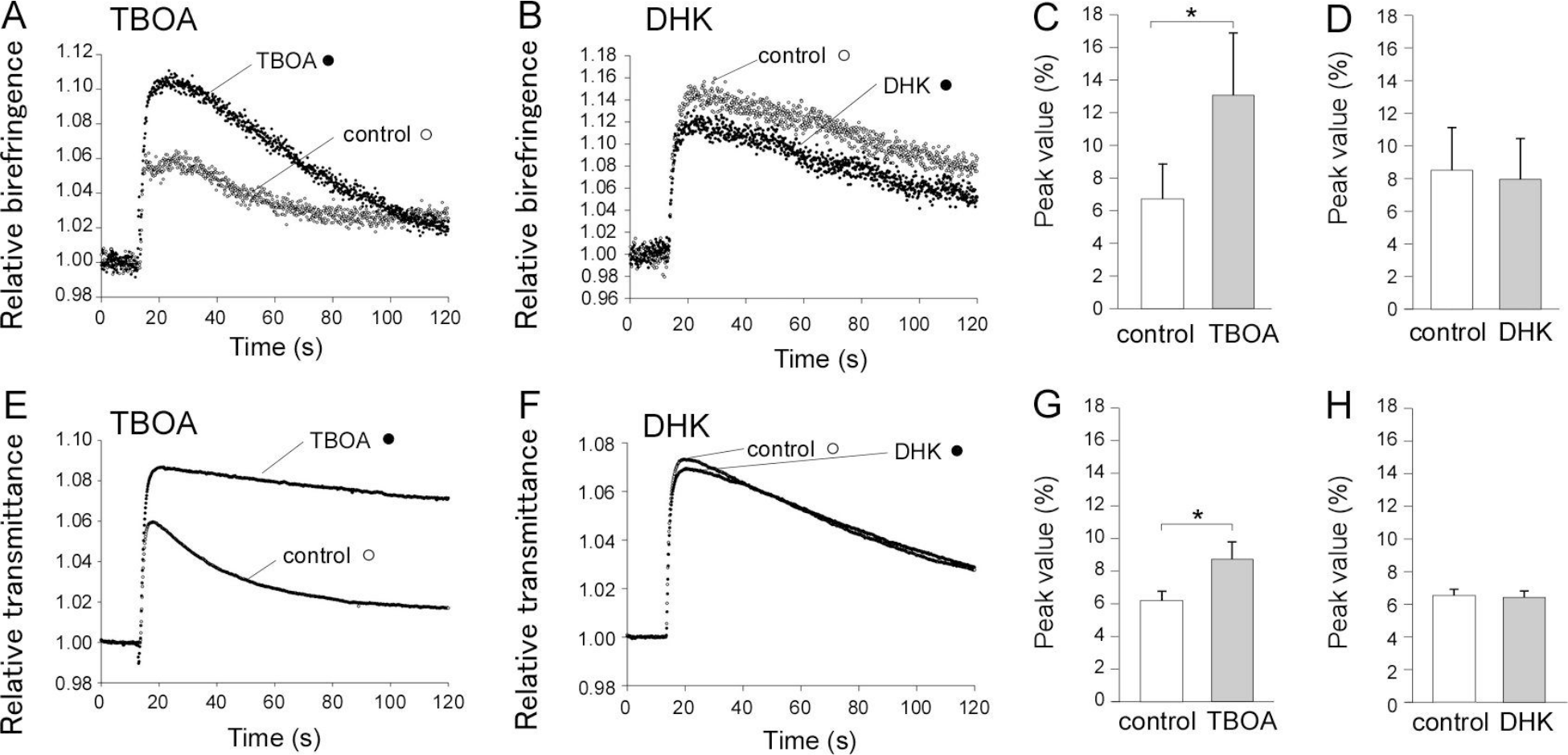
Effects of inhibitors of glutamate transporters on activity dependent relative birefringence change and transmittance change at stratum radiatum of area CA1. (*A*) Representative recordings of relative birefringence changes elicited by electrical stimulation before (*control; open circle*) and after the application of 50µM DL-TBOA (*TBOA; filled circle*). (*B*) Recordings of stimulation-evoked relative birefringence changes before (*control; open circle*) and after the application of 300 µM dihydrokainate (*DHK; filled circle*). (*C*) Summary of the effect of TBOA on the peak amplitude of relative birefringence change. The peak value is expressed as increase in % from the resting birefringence value. 6.74 ± 2.10 % (*control; white bar*), 13.12 ± 3.76 % (*TBOA; gray bar*). Paired t-test, p=0.036 (n=5). (*D*) Summary of the effect of DHK on the peak amplitude of relative birefringence change. 8.42 ± 2.66 % (*control; white bar*), 7.88 ± 2.54 % (*DHK; gray bar*). Paired t-test, p=0.36 (n=5). (*E*) Recordings of stimulation-evoked relative transmittance changes before (*control; open circle*) and after the application of 50 µM DL-TBOA (*TBOA; filled circle*) are superimposed. (*F*) Recordings of stimulation-evoked relative transmittance changes before (*control; open circle*) and after the application of 300µM dihydrokainate (*DHK; filled circle*) are superimposed. (*G*) Summary of the effect of TBOA on the peak amplitude of relative transmittance change. The peak value is expressed as increase in % from the resting transmittance value. 6.20 ± 0.54 % (*control; white bar*), 8.74 ± 1.02 % (*TBOA; gray bar*). Paired t-test, p=0.010 (n=5). (*H*) Summary of the effect of DHK on the peak amplitude of relative transmittance change. 6.52 ± 0.38 % (*control; white bar*), 6.40 ± 0.41 % (*DHK; gray bar*). Paired t-test, p=0.781 (n=5). Data are expressed as mean ± S.E.M. *p<0.050.

We next analyzed the effect of DL-TBOA (Fig. 13E) and DHK (Fig. 13F) on transmittance signal. Similarly to the result in birefringence signal, the peak amplitude of average transmittance was significantly enhanced by DL-TBOA (Fig. 13G, 6.20 ± 0.54 in control, 8.74 ± 1.02 in DL-TBOA, paired t-test, p=0.01, n=5), but not by DHK (Fig. 13H, 6.52 ± 0.38 in control and 6.40 ± 0.41 in DHK, paired *t*-test, p=0.78, n=7).

Our present result cannot discard the possibility of the contribution of GLAST. As previously reported, GLT-1 and EAAC1 show more enriched immunoreactivities than that of GLAST in the mature hippocampus (41, 42). Given that more than 80 % of total glutamate is taken up by GLT-1 and EAAC1 in hippocampus (40 % with GLT-1 and 40 % with EAAC1) (43, 44), the contribution of GLAST is estimated to be less than 20 % of total uptake. The IC_50_ value of DL-TBOA for GLAST is 70 μM (40), which is higher than the concentration that we used for our experiment (50 μM). These reports, together with our present results suggested that EAAC1 could be the major factor that involved in both birefringence and average transmittance changes induced by nerve excitation. The different enhancement effect of DHK and DL-TBOA on birefringence suggests that neuronal glutamate transporter EAAC1 can modify the IOSs by uptaking of spilled over glutamate at perisynaptic regions.

## Discussion

### Polarized light imaging as a label-free method for observing the anatomy and the activity dependent changes of neuronal tissue

One of the novel findings in our observations using advanced polarized light imaging was that the stimulation-evoked IOS changes of the isolated brain involve optical signals raised by anisotropic structural changes in dendrites associated with synaptic transmission. These changes were predominantly detected at the particular locations exhibiting high resting birefringence in the hippocampal slice (Fig. 8E). Traditional IOS (transmittance) imaging did not detect the local difference of stimulation evoked optical responses at the dendritic area of CA1 (Fig. 8D). Thus, the polarization-sensitive IOS reports structural changes in dendritic structure of neurons in response to synaptic transmission that is not detected with the traditional IOS. The involvement of birefringence change in IOS is innovative as the optical anisotropy change suggests additional structural insights into the mechanisms of IOS generation, which by far was optically isotropic change in cellular structures such as cell swelling (5, 6, 9, 13, 29).

### Mapping resting birefringence of acute mouse hippocampal slices

We report for the first time the birefringence map of acute hippocampal slices from mice using LC-PolScope (18, 45). The optical axis of orientation and the retardance value of observed birefringence were correlated with the orientation and the density of dendrites and axons in the hippocampal slice (Fig. 1). Based on the anatomy of hippocampus, Schaffer collateral pathway (axons of CA3 neurons) are aligned perpendicular to the dendrites of CA1 neurons at st. radiatum (30, 46). LC-PolScope demonstrated that the slow axis orientation at st. radiatum was mostly parallel to the net orientation of apical dendrites of CA1 pyramidal neurons, or perpendicular to the net orientation of Schaffer collateral pathway (Fig. 1D). LC- PolScope detects retardance value ranging from sub nm, therefore enables to detect weak birefringence of non-myelinated axons and dendrites in mammalian brains. By imaging individual dendrites in primary cultured neurons, we found two different components of birefringence (Fig. 4). The first component was observed at the cytoplasmic region of dendrites, which slow axis orientation was analyzed as parallel to the longitudinal axis of the dendrites. Microtubules, one of the major components of the cytoskeleton along dendrites, are known to show the slow axis orientation that is parallel to the longitudinal axis of the filaments (47, 48). Observations from our current and past studies suggest that microtubules could be one of the major origins of birefringence signal at the cytoplasm of dendrites. The second component of birefringence was observed at the surface of dendrites, in which the slow axis was perpendicular to the longitudinal axis of dendrites. Previous study by others showed that the slow axis of birefringence originated from lipid membrane was perpendicular to the longitudinal axis of neuronal processes (49–51). Another factor to be considered regarding birefringence source at the surface of dendrites is the edge birefringence (52), which is typically observed at the interface between two media with different refractive indexes. Our result revealed that the absolute retardance value of the cytoplasmic component was twice as high as that of the surface component, therefore the net slow axis orientation of single dendrite was parallel to the dendritic axis.

In our previous study using acute cerebellar slices from chick brains, the slow axis orientation of parallel fibers, which are bundles of non-myelinated axons, was perpendicular to the longitudinal axis. It should be noted that the slow axis orientation of myelinated axons of Purkinje cells was also perpendicular to their length (21), indicating that the bundles of both myelinated and non-myelinated axons show slow axis orientation perpendicular to the length. Olfactory nerves of freshwater pikes were composed of bundles of 4.2 million of non-myelinated axons, in which the slow axis of resting birefringence was perpendicular to the fibers (50, 53). According to these studies, nerve fibers composed of many axons show slow axis orientation primarily originated from the birefringence of surface component.

The diameter of nerve fibers (both axons and dendrites) could be the key factor that determines the net birefringence and the slow axis orientation. In general, dendrites have larger diameter compared with those of axons in the hippocampus. The diameter of proximal dendrites of CA1 pyramidal neurons ranges from 1 to 3 µm, and 0.25 to 1 µm for the distal dendrites. In contrast, the diameter of Schaffer collateral axons is less than 0.2 µm (54, 55). Based on these considerations, the slow axis orientation of birefringence at area CA1 might arise from the constructive contributions of birefringence from thin Shaffer collateral axons in which the slow axis is parallel to the layer of stratum radiatum, and birefringence of dendrites of postsynaptic pyramidal neurons in which the slow axis orientation is parallel to the layer of stratum radiatum. Our basic understanding of the origins of resting birefringence in neuronal processes is generally consistent with the previous reports from by others (49, 50, 53, 56). Resting birefringence of neuronal tissue can be preserved even after chemical fixation of samples as long as both the cytoplasmic and the lipid membrane components are well preserved (56). Polarized light imaging approach can be applied to map 3D wiring of fiber tracts of brain preparations from humans and primates without staining the tissue (57–59).

### Origins of birefringence changes in mouse hippocampal slices that associate with neuronal activity

The instantaneous PolScope enabled to detect changes of polarization-sensitive IOS (relative birefringence) at area CA1 of hippocampus followed by electrical stimulation at Schaffer collaterals. The spreading and the temporal change of the polarization-sensitive IOS were essentially the same as those observed with traditional IOS, except for the decay time of relative birefringence was faster than the traditional IOS (Fig. 7). Our results using blockers of AMPA-type and NMDA-type glutamate receptors indicated that approximately 70% of synaptically induced birefringence change was due to postsynaptic factors. The contributions of postsynaptic neurons to the generation of pol-IOS changes are similar to the results that had been reported using traditional IOS imaging (5, 9). These observations suggested that the mechanisms of birefringence changes evoked by neuronal excitation could be involved in the mechanisms that generate traditional IOS. MacVicer et al. proposed that the main cause of IOS increase associated with synaptic transmission in brain slices is due to postsynaptic cell swelling induced by water-influx through the membrane (5, 6, 29). As the water influx induce the reduction of refractive index of cytoplasm, the water influx after synaptic transmission may lower the edge birefringence between the cytoplasm of dendrites in CA1 pyramidal neurons and the surrounding ACSF. However, as we have noted before, in some patched areas of dendritic region of CA1 where the resting birefringence values were higher than surrounding areas, the stimulation evoked pol-IOS changes (birefringence changes) were more prominent than raditional IOS (transmittance changes) (Fig. 8). These observations might require other reasoning that involves resting birefringence in the mechanisms of pol-IOS generation.

One of the other possibilities of the birefringence change is the structural reorganization of postsynapses that is associated with synaptic input. In particular, the modifications of spines in their shape, volume and number in activity dependent manner (60, 61), can be the origins of birefringence changes. Since actin filaments generate birefringence signals (20, 28), changes in F-actin architectures could contribute to overall birefringence change in dendritic area at CA1 followed by synaptic transmission. FRET imaging demonstrated that the tetanic stimulation enhanced the polymerization of actin that enlarged spine heads (62).

Our pharmacological data suggests that 70 % of birefringence signal observed at area CA1 involves the activation of glutamate receptors (Fig.12). Previous studies support excitatory synaptic transmission alters postsynaptic organization of actin filaments via AMPA and NMDA receptor activation (63). As described above, architectural change of actin filaments in spines and dendrites through postsynaptic AMPA and/or NMDA receptors activation can be one of the sources of birefringence change observed in our polarized light imaging.

The rest of 30 % of the birefringence change could be presynaptic origin and/or glial cell origins. This can involve membrane dynamics at presynaptic terminals, in particular vesicular endocytosis followed by excitatory synaptic transmission. Recent study demonstrates the time course of clathrin-mediated vesicular endocytosis recorded at presynaptic terminal after repetitive action potential was _slow_ 26 sec (64). Our data showed that the decay time constant of birefringence changes was 30.3 ± 2.35 sec (Fig. 10), which was close to the time constant for internalization of synaptic vesicles. This evidence supports that the presynaptic membrane dynamics could be involved in the mechanisms underlying birefringence change associated with neuronal excitation.

Since microtubules are one of the major birefringence sources in living cells (47, 48), and identified as a key molecule to modify vesicular dynamics at presynaptic terminals (65), microtubules might be responsible on the mechanisms underlying presynaptic origin of birefringence change. Recently, capacitance measurement directly applied to excitatory presynaptic terminals revealed the assembly of microtubules modulate the speed of vesicular endocytosis (66). Change in the number of synaptic vesicles may cause birefringence change as the ratio of isotropic, spherical membranes and anisotropic, tubular or button-shape membranes determines the net birefringence of synapses. This is consistent with our data that relative birefringence increased followed by repetitive neuronal stimulation, as the decrease in the number of synaptic vesicles in presynaptic terminals during the synaptic transmission. However, dynamics of microtubules that correlate with neuronal birefringence change has been left for further studies.

